# Diet-Induced Vitamin D Deficiency Results in Reduced Skeletal Muscle Mitochondrial Respiration in C57BL/6J Mice

**DOI:** 10.1101/2020.05.15.098087

**Authors:** Stephen P. Ashcroft, Gareth Fletcher, Ashleigh M. Philp, Philip J. Atherton, Andrew Philp

## Abstract

Vitamin D deficiency is known to be associated with symptoms of skeletal muscle myopathy including muscle weakness and fatigue. Recently, vitamin D related metabolites have been linked to the maintenance of mitochondrial function within skeletal muscle. However, current evidence is limited to *in vitro* models and the effects of diet-induced vitamin D deficiency upon skeletal muscle mitochondrial function *in vivo* have received little attention. In order to examine the role of vitamin D in the maintenance of mitochondrial function *in vivo*, we utilised an established model of diet-induced vitamin D deficiency in C57BL/6J mice. Mice were fed either a control (2,200 IU/kg) or a vitamin D deplete (0 IU/kg) diet for periods of 1-, 2- and 3-months. Skeletal muscle mitochondrial function and ADP sensitivity were assessed via high-resolution respirometry and mitochondrial protein content via immunoblotting. As a result of 3-month of diet-induced vitamin D deficiency, respiration supported via CI+II_P_ and ETC were 35% and 37% lower when compared to vitamin D replete mice (*P* < 0.05). Despite functional alterations, the protein expression of electron transfer chain subunits remained unchanged in response to dietary intervention (*P* > 0.05). In conclusion, we report that 3-months of diet-induced vitamin D deficiency reduced skeletal muscle mitochondrial function in C57BL/6J mice. Our data, when combined with previous *in vitro* observations, suggests that vitamin D mediated regulation of mitochondrial function may underlie the exacerbated muscle fatigue and performance deficits observed during vitamin D deficiency.

## Introduction

Vitamin D deficiency, characterised by serum 25(OH)D levels of <50 nmol.L^-1^, remains prevalent across both Europe and the USA [1 2]. Although the classical actions of vitamin D within the maintenance of bone health are well established [3-5], a number of non-classical actions have recently been identified including the maintenance of skeletal muscle function [6].

Within human populations, multiple observational studies have reported a positive association between serum 25(OH)D, skeletal muscle strength and lower extremity function in older individuals [7-9]. Furthermore, the supplementation of vitamin D has also been reported to increase muscle strength within this population [10 11]. Despite these associations, studies of this design are unable to infer causality. In addition, isolating the effects of vitamin D status within older populations is often difficult given individuals may suffer from a number of pre-existing conditions that may interfere with vitamin D status [12]. These difficulties highlight the importance of model systems that allow for the manipulation and isolation of vitamin D status in order to study the precise role of vitamin D within skeletal muscle.

In order to study the impact of vitamin D deficiency on skeletal muscle function, a number of animal models have been utilised. A dysregulation of vitamin D status can be achieved via dietary means [13-16], a reduction in sunlight exposure [13] or by the administration of ethane 1-hydroxy-1, 1-diphosphonate which blocks the production of 1α,25-dihydroxyvitamin D_3_ (1α,25(OH)_2_D_3_) [16]. Diet-induced vitamin D deficiency has been shown to result in symptoms of skeletal muscle myopathy including impaired contraction kinetics, skeletal muscle weakness, as well as decreases in muscle force in both chicks and rats [13 14 17]. In order to isolate the effects of vitamin D alone and offset the observed hypocalcemia and hypophosphatemia that are associated with the induction of vitamin D deficiency [17], diets with increased calcium and phosphate have been utilised [15]. However, despite the administration of this rescue diet, mice still display reduced grip strength and an increase in *Myostatin* gene expression [15], a known negative regulator of muscle mass [18]. Similarly, mice fed this diet chronically (8-12 months) show similar impairments in physical performance including; reduced grip endurance, sprint speed and stride length [19].

The observed impairments in physical performance with vitamin D deficiency may be linked to skeletal muscle mitochondrial function [20 21]. *In vitro*, vitamin D related metabolites are able to increase mitochondrial function in both immortalised and primary skeletal muscle cell lines [22-25]. Furthermore, we recently observed significant impairments in mitochondrial function in Vitamin D Receptor (VDR) loss- of-function C2C12 myoblasts [26]. In humans, the supplementation of vitamin D within a cohort of severely deficient individuals resulted in a reduced phosphocreatine (PCr) recovery time, as measured non-invasively by 31-phosphorous magnetic resonance spectroscopy (31-P MRS) [20]. Whilst skeletal muscle mitochondrial content seems to remain unchanged following diet-induced vitamin D deficiency in mice [19], the functional characteristics of the mitochondria remain largely underexplored. Therefore, we aimed to determine the effects of diet-induced vitamin D deficiency upon skeletal muscle mitochondrial function in C57BL/6J mice.

## Methods

### Ethical Approval

Ethical approval was granted by the Garvan Institute and St. Vincent’s Hospital Animal Experimentation Ethics Committee (approval number 18/19), fulfiling the requirements of the NHMRC and the NSW State Government, Australia. All animal handling was carried out by trained personnel and all procedures were carried out according to the Australian code of practice for the care and use of animals for scientific purposes 8^th^ edition [27]. Male C57BL/6JAusb mice were received at 10-weeks of age and housed communally in a temperature controlled environment (22 ± 0.5°C) with a 12 h light-dark cycle.

### Composition of Diet

Following 1-week acclimation in which mice were fed a standard chow diet, mice were placed on either a vitamin D-control diet (SF085-034, Speciality Feeds, Glen Forest, NSW) or a vitamin D-deplete diet (SF085-003, Speciality Feeds, Glen Forest, NSW) for periods of 1- (n = 10/group), 2- (n = 6/group) or 3-months (n = 6/group). The vitamin D deplete contains no vitamin D (cholecalciferol 0 IU/kg) but increased calcium (2%) and phosphorous (1.2%) in order to maintain normal mineral homeostasis. Previously, this dietary intervention has been shown to successfully induce vitamin D deficiency following 1-month of dietary intervention [15]. The vitamin D control diet contains vitamin D (cholecalciferol 2,200 IU/kg), calcium (1%) and phosphorous (0.7%).

### Assessment of Food Intake

Food intake was assessed on a monthly basis at 1-, 2- and 3-months of dietary intervention. The weight of the food within the cage was recorded and subsequently re-weighed following a period of 24 h. The amount of food consumed was then divided by the number of mice within the cage and reported as food intake in grams per mouse.

### Assessment of Body Composition

Body weight was obtained on a weekly basis throughout the dietary intervention periods. In addition, prior to each measurement of body composition mice were briefly weighed. Body composition was assessed upon arrival (10-weeks of age) and then following 1-, 2- and 3-months of dietary intervention using the EchoMRI (EchoMRI LLC, Houston, USA).

### Tissue Collection

Tissue collections were completed following 1-, 2- and 3-months of dietary intervention. All samples were excised from fasted (2 h) mice following isoflurane (5%) anesthetization. Following collection, a blood sample was taken via cardiac puncture and animal terminated via cervical dislocation. All tissues were rinsed in sterile saline, blotted dry, weighed, and frozen in liquid nitrogen. A small portion (∼20 mg) of the red gastrocnemius was removed before freezing and used for high-resolution respirometry. All further tissue samples were stored at -80°C for subsequent analysis.

### Tissue Processing

Small portions of red gastrocnemius muscle (∼20 mg) were removed and placed in ice-cold BIOPS buffer (2.77 mM CaK_2_EGTA, 7.23 mM K_2_EGTA, 5.77 mM Na_2_ATP, 6.56 mM MgCl_2_-6H_2_O, 20 mM Taurine, 15 mM Na_2_Phosphocreatine, 20 mM Imidazole, 0.5 mM Dithiothreitol, 50 mM MES Hydrate, pH 7.1, 290 mOsm). Blood samples were allowed to coagulate at room temperature for 10 minutes before being placed on ice. Blood samples were then centrifuged at 14,000 *g* for 10 minutes and the resulting supernatant was removed and stored at -80°C for further analysis.

### Analysis of Serum Calcium

Serum calcium was measured using a Calcium Detection Assay kit (Abcam, Cambridge, UK). Serum samples were diluted 1:10 and manufacturers instructions were followed. The assay plate was read at 575 nm using a CLARIOstar microplate reader (BMG Labtech, Victoria, Australia). Serum calcium concentrations are reported in mM.

### High-Resolution Respirometry

High-resolution respirometry was conducted in MiR05 (2 ml) with the addition of blebbistatin (25 μM) using the OROBOROS Oxygraph-2K (Oroboros Instruments, Corp., Innsbruck, AT) with stirring at 750 rpm at 37°C. Oxygen within the chamber was maintained between 150-220 μM for each experiment. Prior to the addition of the fibre bundles to the chamber, bundles were blotted dry and weighed. Bundles totalling 2.5-5.0 mg were added to each chamber. Firstly, pyruvate (10 mM) and malate (2 mM) were added in assessment of complex I related leak (CI_L_). ADP was then titrated in step-wise increments (100-6000 μM) followed by the addition of glutamate (10 mM) to assess phosphorylating respiration (CI_P_). The addition of succinate (10 mM) followed to assess respiration support via complex II (CI+II_P_). Cytochrome c (cyt c) (10 μM) was added in order to check outer mitochondrial membrane integrity. The partial loss of cyt c during fibre preparation may limit respiration however, no fibre preparation exhibited an increase of >10%. Carbonyl cyanide 3-chlorophenylhydrazone (CCCP) was titrated in a step-wise manner (0.5 to 2.5 μM) until the maximal capacity of the electron transport chain (ETC) was reached. Finally, antimycin A (2.5 μM) was injected in order to determine non-mitochondrial oxygen consumption.

The apparent K_m_ for ADP was determined through the Michaelis-Menten enzyme kinetics – fitting model (Y = Vmax*X/(K_m_ + X)), where X = (free ADP; ADP_f_), using Prism (GraphPad Software, Inc., La Jolla, CA) as previously described [28]. Flux control ratios (FCR) was calculated by setting CCCP stimulated respiration as 1 and antimycin A respiration as 0.

### Immunoblotting

Gastrocnemius samples were powdered on dry ice using a Cellcrusher™ tissue pulverizer (Cellcrusher Ltd, Cork, Ireland) and homogenized via shaking in a FastPrep 24 5G (MP Biochemicals, Santa Ana, California, USA) at 6.0 m·s^-1^ for 80 s in a 10-fold mass of ice-cold sucrose lysis buffer (50 mM Tris pH 7.5; 270 mM sucrose; 1 mM EDTA; 1 mM EGTA; 1% Triton X-100; 50 mM sodium fluoride; 5 mM sodium pyrophosphate decahydrate; 25 mM beta-glytcerolphosphate). Inhibitors were added fresh on the day of use and included 1 cOmplete™ protease inhibitor cocktail EDTA free tablet (Roche, Basel, Switzerland) and Phosphatase Inhibitor Cocktail 3 both purchased from Sigma-Aldrich (Sigma-Aldrich, NSW, Australia).

Samples were then centrifuged for 10 min at 8,000 *g* at 4°C to remove any insoluble material. Protein concentrations were determined using the DC protein assay as per manufacturer’s instructions (Bio-Rad, NSW, Australia). An equal volume of protein (30 μg) was separated by SDS-PAGE on 12.5% gels at a constant current of 23 mA per gel for ∼60 minutes. Proteins were then transferred on to BioTrace NT nitrocellulose membranes (Pall Life Sciences, Pensacola, Florida, USA) using a wet transfer system at 100 V for 1 h. Membranes were then stained in Ponceau S (Sigma-Aldrich, NSW, Australia) and imaged to check for even loading and transfer. Membranes were then blocked for 1 h in 3% dry-milk in tris-buffered saline with tween (TBS-T). Membranes were incubated overnight in primary antibodies at 4°C. Following primary antibody incubation, membranes were washed three times in TBS-T and subsequently incubated in the appropriate horseradish peroxidase-conjugated secondary antibody at room temperature for 1 h. Membranes were again washed three times in TBS-T prior to imaging. Images were captured using the ChemiDoc (Bio-Rad, NSW, Australia) and quantified using ImageJ.

### Antibodies

MitoProfile OXPHOS antibody cocktail (110413) was from Abcam and used at a concentration of 1:1,000. Anti-mouse (7076) secondary antibody was used at a concentration of 1:10,000 in TBS-T and was from Cell Signaling Technology.

### Statistical Analysis

Statistical analysis was performed using Prism version 7 (GraphPad Software Incorporated, La Jolla, CA, USA). Differences between 1-, 2- and 3-month vitamin D replete and deplete mice were determined by two-way ANOVA with Bonferroni correction for multiple comparisons. Differences between vitamin D deplete and replete mice in mitochondrial respiration in response to ADP titration were determined by multiple t-test. All values are presented as mean ± SD. Statistical significance was set at *P* < 0.05.

## Results

### Body Composition Following Diet-Induced Vitamin D Deficiency

Body weight increased across time when mice were compared at the 1-, 2- and 3-month time-points (*P* < 0.001) (Fig. 1A). However, no differences in body weight were observed when comparing vitamin D replete and deplete mice (*P* > 0.05) (Fig. 1A). Further assessment of body composition revealed no differences in absolute lean mass (*P* > 0.05) (Fig. 1C) however, when expressed as a percentage of body weight, lean mass was lower when compared across dietary intervention time-points (*P* < 0.001) (Fig. 1D). Further analysis revealed that lean mass as a percentage of body weight was 16% lower at 3-months when vitamin D deplete mice were compared with the 1-month vitamin D deplete (*P* < 0.001) (Fig. 1D). Although this is indicative of a loss of lean mass with vitamin D deficiency, this effect was potentially driven by a higher lean mass as a percentage of body weight at the 1-month time-point in vitamin D deplete mice (*P* = 0.039) (Fig. 1D). Both absolute and percentage of body weight fat mass increased across time in both dietary groups (*P* < 0.001) (Fig. 1E-F). Furthermore, fat mass as a percentage of body weight was 60% higher in the vitamin D replete mice when compared with vitamin D deplete at the 1-month time point (*P* = 0.044) (Fig. 1F). Despite differences in body composition, no difference in food intake was observed between time points (*P* > 0.05) and groups (*P* > 0.05) (Fig. 1B).

**Figure 1.**
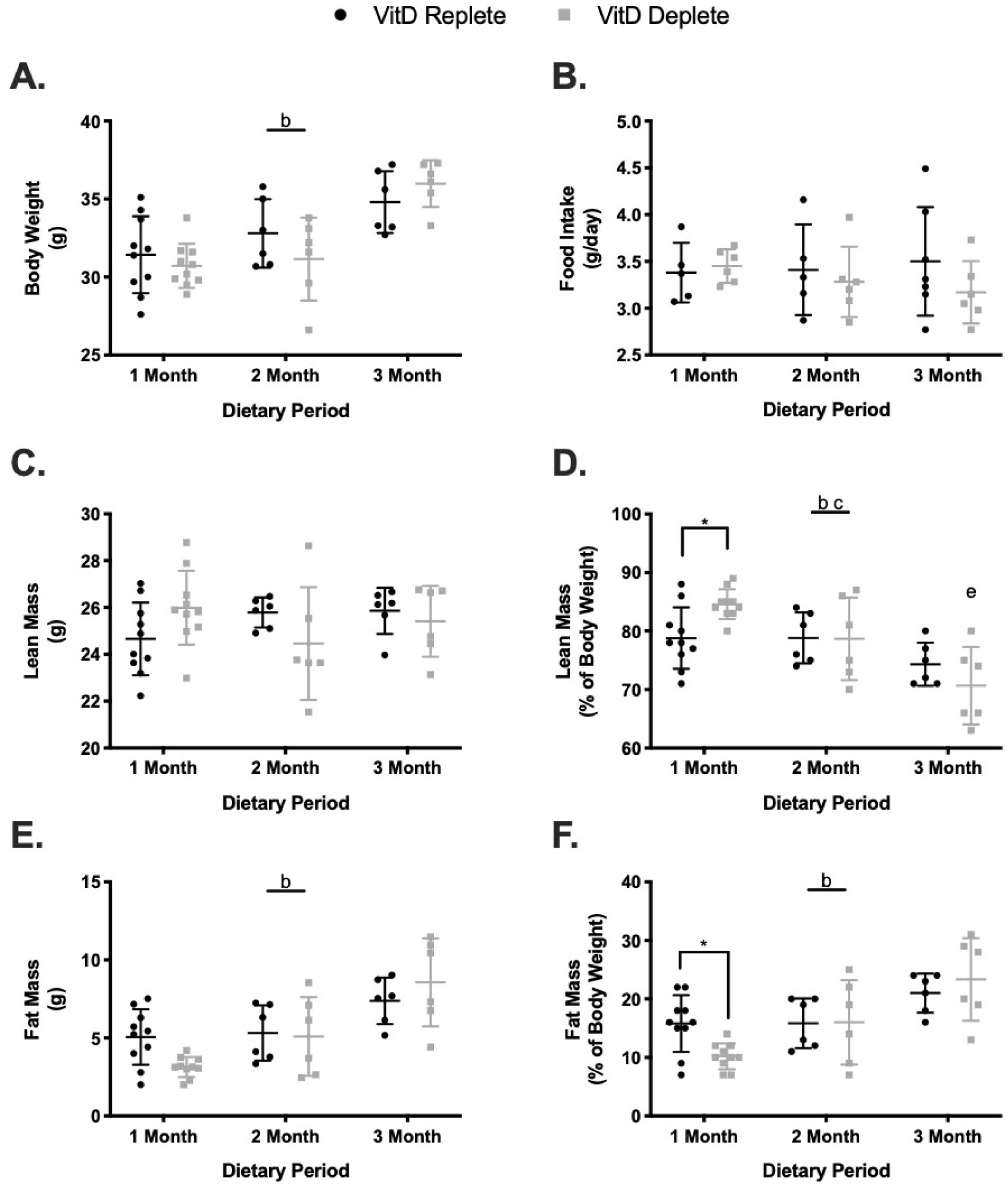
Assessment of body composition and food intake in vitamin D replete and deplete C57BL/6J mice. A, increase in body weight across dietary period with no differences between differing vitamin D diets (n = 6-10/group). B, no difference in food intake was observed following dietary intervention (n = 5-6 cages/group). C-D, absolute lean mass remained unchanged whilst lean mass as a percentage of body weight was significantly lower in the 3-month vitamin D deplete when compared to the 1-month (n = 6-10/group). E-F, Absolute and percentage of body weight fat mass increased across the dietary period irrespective of dietary intervention. Data mean ± SD. ^b^ Main effect for time (*P* < 0.05), ^c^ Main effect group x time interaction (*P* < 0.05), * (*P* < 0.05).

### Skeletal Muscle Mass

Given the alterations in total lean mass, we determined whether individual skeletal muscle mass was effected by diet-induced Vitamin D deficiency. Crude analysis of skeletal muscle wet weight revealed no differences in the mass of the gastrocnemius in response to either dietary intervention (*P* = 0.408) or time point (*P* = 0.103) (Table 1). Overall, the mass of the quadriceps increased over time (*P* = 0.004) however, this was not changed by dietary intervention (*P* = 0.951) (Table 1). Collectively, triceps mass was higher when vitamin D replete mice were compared with deplete (*P* = 0.041) although, post-hoc analysis revealed no difference between groups at individual time points (*P* > 0.05) (Table 1).

**Table 1.** Assessment of skeletal muscle wet weight following the induction of diet-induced vitamin D deficiency (n = 6-10/group). Data mean ± SD.

|  | Dietary Period (Months) |  |  | *P* |  |
| --- | --- | --- | --- | --- | --- |
|  | 1 | 2 | 3 | *VitD* | *Time* |
| **Gastrocnemius** |  |  |  |  |  |
| (mg) |  |  |  |  |  |
| Replete | 143 ± 14 | 166 ± 25 | 161 ± 14 |  |  |
| Deplete | 152 ± 16 | 149 ± 12 | 156 ± 15 | 0.408 | 0.103 |
| **Quadriceps** |  |  |  |  |  |
| (mg) |  |  |  |  |  |
| Replete | 160 ± 14 | 189 ± 12 | 182 ± 20 |  |  |
| Deplete | 169 ± 18 | 178 ± 22 | 184 ± 19 | 0.951 | 0.004 |
| **Tricep** |  |  |  |  |  |
| (mg) |  |  |  |  |  |
| Replete | 111 ± 20 | 113 ± 14 | 109 ± 10 |  |  |
| Deplete | 106 ± 9 | 96 ± 19 | 101 ± 13 | 0.041 | 0.695 |

### Serum Calcium

Similar to previous reports [15], we observed no change in serum calcium irrespective of dietary group or time point (*P* > 0.05) (Fig. 2A).

**Figure 2.**
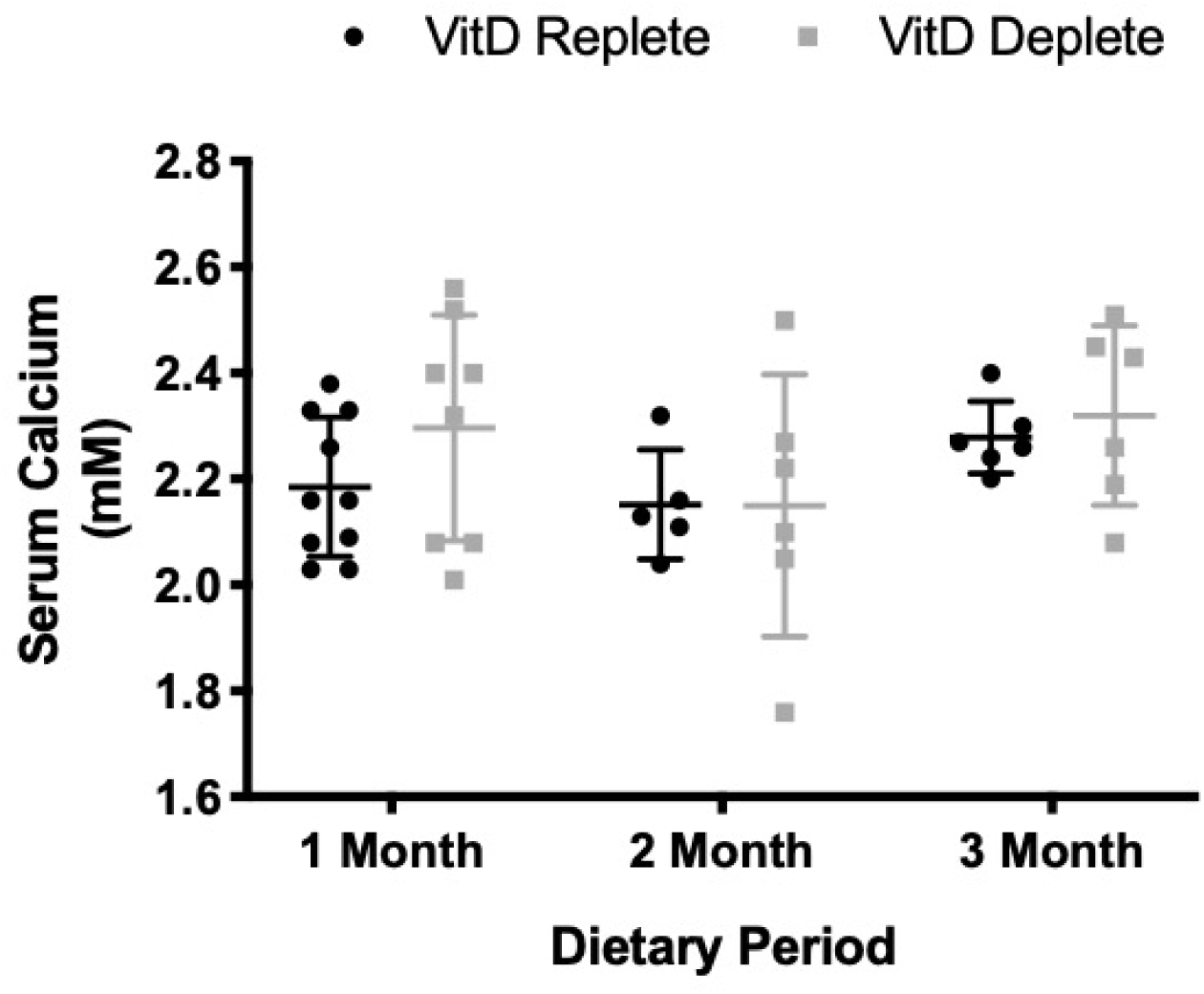
Serum calcium in response to the induction of diet-induced vitamin D deficiency (n = 5-10/group). Data mean ± SD.

### Mitochondrial Function

In response to both pyruvate and malate alone, we observed no change in CI_L_ (*P* > 0.05) (Fig. 3A). However, following the addition of ADP, CI_P_ respiration increased across the 1, 2 and 3-month time points (*P* = 0.048) (Fig. 3B). Furthermore, respiration was 85% and 96% higher at the 2- (*P* = 0.015) and 3-month (*P* = 0.006) time-points respectively when compared with 1-month in vitamin D replete mice (Fig. 3B). Similarly, higher respiratory capacities during CI+II_P_ respiration and the maximal capacity of the ETC were observed in the 2- and 3-month vitamin D replete mice when compared with the 1-month (*P* < 0.05) (Fig. 3C-D). In addition, 3-months of diet-induced vitamin D deficiency resulted in 35% and 37% lower respiratory rates during CI+II_P_ (*P* = 0.035) and maximal ETC capacity (*P* = 0.015) when compared to vitamin D deplete mice at the same time-point (Fig. 3C-D). We also analysed to above data as a flux control ratio which provides a method for internal normalisation [31 32]. Despite the observed changes reported above, flux control ratios revealed no differences in mitochondrial respiration supported via complex I alone (*P* > 0.05) or complex I and II in combination (*P* > 0.05) (Fig. 3E-F).

**Figure 3.**
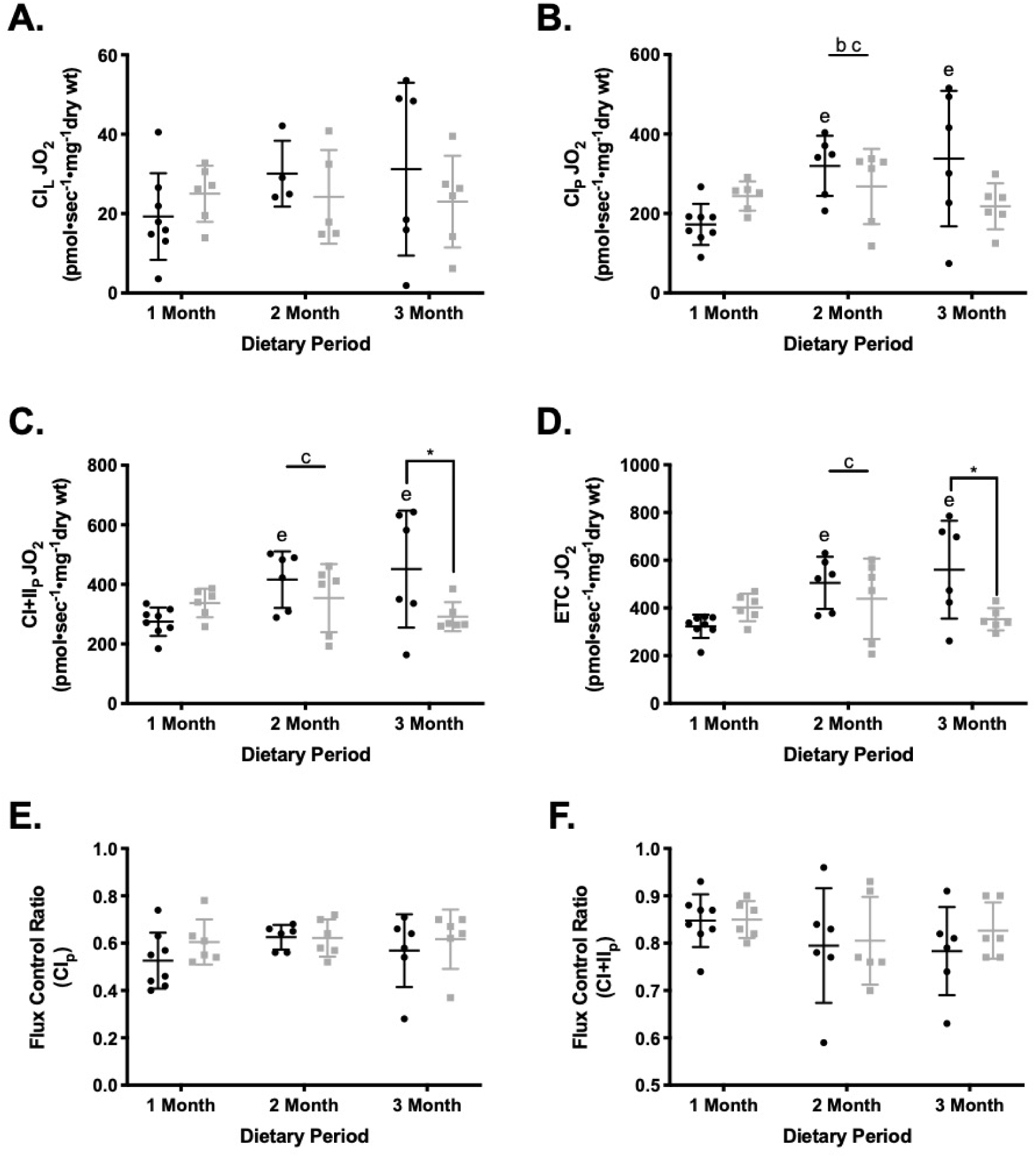
Assessment of skeletal muscle mitochondrial respiration in response to diet-induced vitamin D deficiency. A, CI_L_ remains unchanged irrespective of dietary intervention (n = 4-8/group). B, Respiration supported via CI_P_ is higher in the 2- and 3-month vitamin D replete mice when compared with the 1-month (n = 5-8/group). C-D, Respiration supported via CI+II_P_ and the maximal capacity of the ETC is increased in the vitamin D replete mice and higher at the 3-month time point when compared to vitamin D deplete (n = 5-8/group). E-F, alteration in absolute rates of respiration are diminished when internally normalised (n = 5-8/group). Data mean ± SD. ^b^ Main effect for time (*P* < 0.05), ^c^ Main effect group x time interaction (*P* < 0.05), * (*P* < 0.05).

### ADP Sensitivity

Commonly, the assessment of mitochondrial function is performed under saturating concentrations of ADP [31 32] which may not be biologically relevant. Therefore, we assessed mitochondrial function in response to a titration of ADP from biologically relevant to saturating concentrations [28 33]. We observed no differences in mitochondrial respiration throughout the titration of ADP between vitamin D replete and deplete mice (*P* > 0.05) (Fig. 4A-C). Furthermore, no differences were observed in the apparent K_m_ for ADP in response to either dietary intervention (*P* > 0.05) or time point (*P* > 0.05) (Fig. 4D).

**Figure 4.**
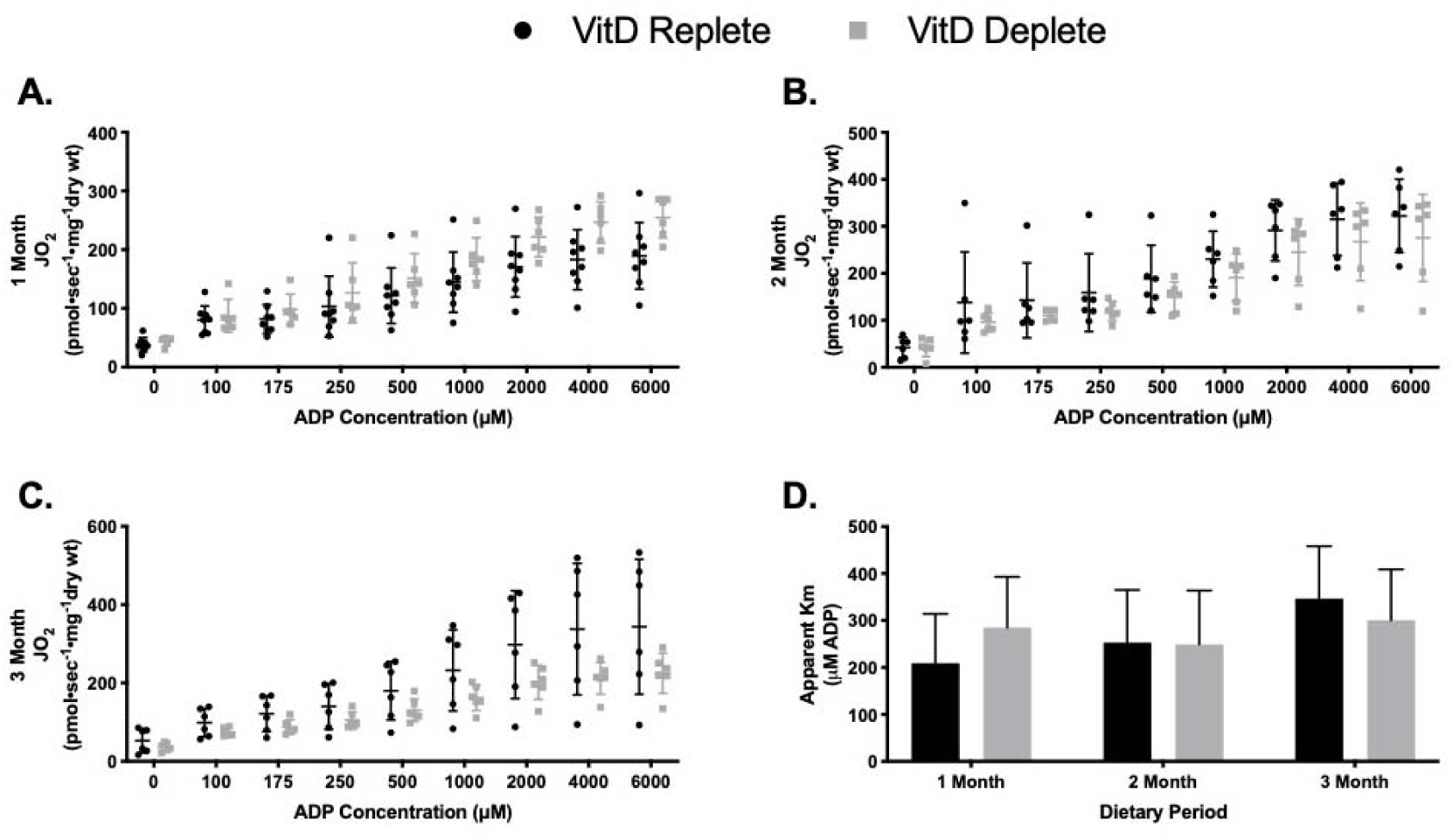
Assessment of skeletal muscle mitochondrial ADP sensitivity in response to diet-induced vitamin D deficiency. A-C, no change in respiratory capacity in response to the titration of ADP following 1-, 2- and 3-month of vitamin D deficiency (n = 5-8/group). D, no change in the apparent K_m_ for ADP in response to 1-, 2- and 3-month of vitamin D deficiency (n = 5-8/group). Data mean ± SD.

### Mitochondrial Protein Content

Finally, given the observed decrements in mitochondrial function associated with vitamin D deficiency, we sort to assess mitochondrial protein content following diet-induced vitamin D deficiency. We observed no changes in complex I (NDUFB8), complex II (SDHB) and complex IV (UQCRC2) protein content when compared irrespective of time point or vitamin D diet (*P* > 0.05) (Fig. 5A-B, D). Interestingly, both complex III (MTCO1) and complex V (ATP5A) were decreased across time (*P* < 0.05) however, there were no differences between dietary groups (*P* > 0.05) (Fig. 5C & E).

**Figure 5.**
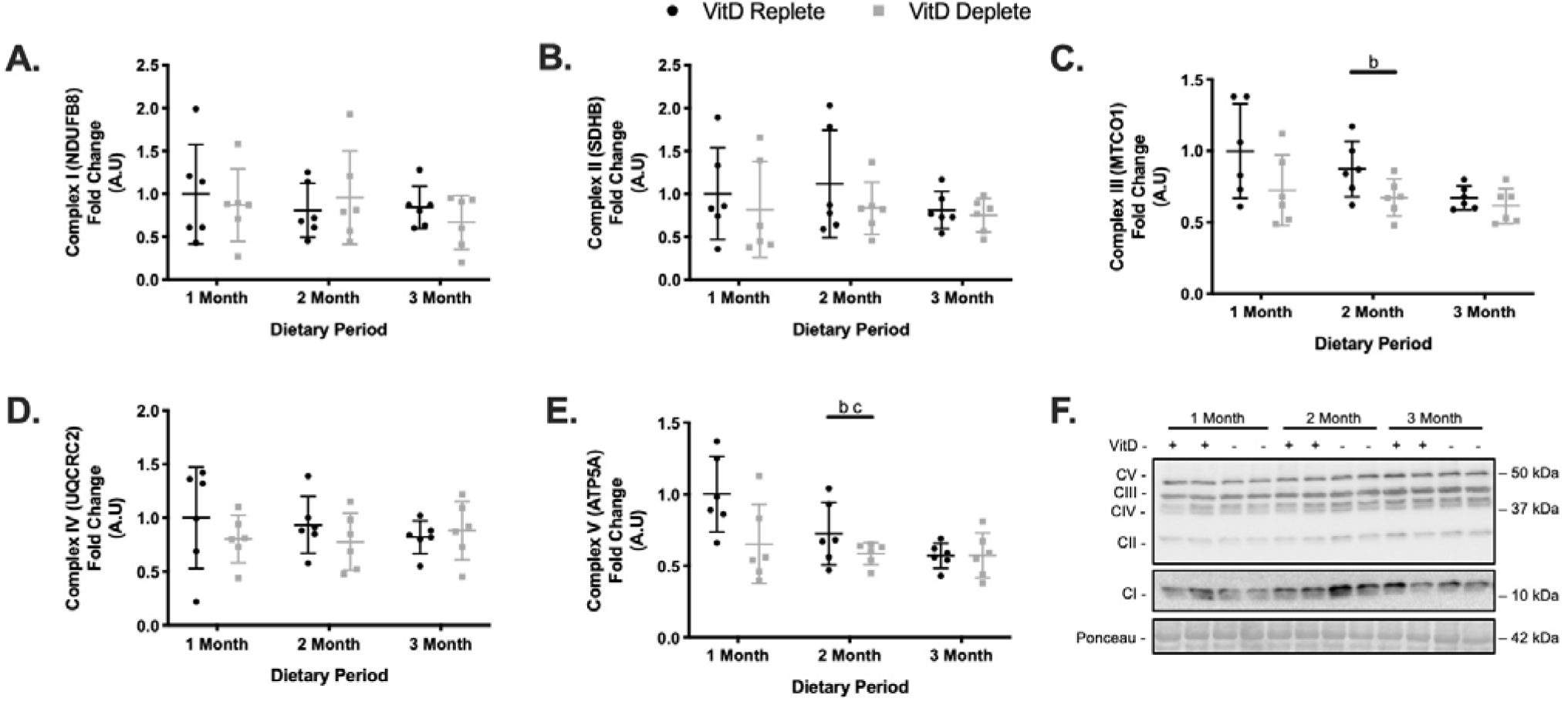
Assessment of skeletal muscle mitochondrial protein content following diet-induced vitamin D deficiency. A-B and D, no changes in expression levels of mitochondrial complex I (NDUFB8), complex II (SDHB) and complex IV (UQCRC2) following diet-induced vitamin D deficiency (n=6/group). C and E, lower expression levels of mitochondrial complex III (MTCO1) and complex V (ATP5A) across the dietary time period with no differences between vitamin D replete and deplete mice (n = 6/group). Data mean ± SD. ^b^ Main effect for time (*P* < 0.05), ^c^ Main effect group x time interaction (*P* < 0.05).

## Discussion

Vitamin D deficiency has been linked to reductions in muscle function, however the specific role of Vitamin D on mitochondrial function in less established. Therefore, the aim of the present study was to directly examine the effects of diet-induced vitamin D deficiency upon skeletal muscle mitochondrial function in C57BL/6J mice. Utilising the current gold standard method to assess mitochondrial function in permeabilised skeletal muscle fibres [29 30], we report that 3-months of diet-induced vitamin D deficiency reduces mitochondrial respiration supported via CI+II_P_ and the maximal capacity of the ETC (Fig. 3C-D). Interestingly, despite the functional changes, we observed no differences in mitochondrial protein content following the induction of diet-induced vitamin D deficiency (Fig. 5A-F). In addition, 1-month of diet-induced vitamin D deficiency resulted in an increase in lean mass (Fig. 1D) and a decrease in fat mass (Fig. 1F) as a percentage of body weight, although these effects were transient as they did not manifest over 2- and 3-months of dietary intervention. Furthermore, diet-induced vitamin D deficiency resulted in a decrease in lean mass as a percentage of body weight across the 3-month time period (Fig. 1D). Despite this, no changes in body weight, lean mass, fat mass or food intake were apparent when comparing vitamin D replete to the deplete group following 3-months of dietary intervention (Fig. 1A-F).

The ability of vitamin D and related metabolites to increase skeletal muscle mitochondrial function in immortalised and primary cell lines has been well established [22-25]. Despite this, there is little evidence for the effects of vitamin D status upon skeletal muscle mitochondrial function *in vivo*. To date, just one study has examined skeletal muscle mitochondrial function, as measured non-invasively via 31-P MRS, in a cohort of severely deficient patients [20]. The authors reported a decrease in PCr recovery time following supplementation with vitamin D, indicative of increased oxidative phosphorylation [20]. However the study followed an open label design, making the extrapolation of this data unclear. In order to directly assess the effects of vitamin D status upon skeletal muscle mitochondrial function, we utilised a mouse model of diet-induced vitamin D deficiency. This model allows for the manipulation of vitamin D status without altering mineral homeostasis [15]. Following 3-months of diet-induced vitamin D deficiency, we report that respiration supported via CI+II_P_ and the maximal capacity of the ETC is lower when compared to vitamin D replete mice at the same time-point. Reduced rates of respiration following 3-months of diet-induced vitamin D deficiency were mediated by increased respiration in the vitamin D replete cohort as opposed to a reduction in vitamin D deplete mice. Following internal normalisation, the above alterations in mitochondrial function were no longer apparent, suggesting the changes observed were due to alterations in mitochondrial quantity as opposed to quality. Therefore, in order to ascertain whether these increases were mediated by an increase in mitochondrial protein abundance, we assessed the protein content of ETC subunits I-V. Despite functional impairments, we observed no differences in markers of mitochondrial protein content within skeletal muscle following the induction of vitamin D deficiency. Therefore, this suggests that vitamin D deficiency alters mitochondrial function independently of mitochondrial content. Similar observations have been made *in vitro* following either the treatment of skeletal muscle cell lines with vitamin D metabolites or the knock-down of the VDR [22 26]. In addition, within the skeletal muscle of the recently developed skeletal muscle muscle-specific VDR-KD mouse, mitochondrial protein content remained unchanged [34]. The further examination of mitochondrial function in this mouse model would help to determine whether the effects of vitamin D deficiency upon skeletal muscle mitochondrial are a direct result of vitamin D related signalling.

In addition, we also sought to determine the effects of diet-induced vitamin D deficiency upon body composition within C57BL/6J mice. Previously, no differences in body weight and lean mass were observed following 12-months of diet-induced deficiency in male C57BL/6J mice [19]. In female mice however, 12-months of diet-induced vitamin D deficiency resulted in reductions in body weight, lean mass and fat mass [35]. Similar to previous reports in male mice, we observed no differences between the vitamin D replete and deplete groups in body weight, lean mass, fat mass or food intake at the 3-month time point. We did however observe a reduction in lean mass as a percentage of body weight from 1- to 3-months in vitamin D deplete mice whereas replete mice remained stable over the same time period. This may in part be driven by the fact we also observed an increase in lean mass as a percentage of body weight following 1-month of diet-induced deficiency. Despite minimal differences in body composition, previous data suggests that vitamin D deficiency impairs physical function in mice [15 19]. Therefore, functional measures of muscle performance may be more relevant to assess the effects of vitamin D deficiency within skeletal muscle. However, it should also be noted that those with serum concentrations of 25(OH)D <25 nmol.L^-1^ are at a greater risk of developing sarcopenia [36]. Given the effects of vitamin D deficiency and supplementation seem to be most potent in older individuals, the induction of vitamin D deficiency in aging mouse models may reveal more potent effects on skeletal muscle mass.

In conclusion, we report that mitochondrial function (CI+II_P_ and ETC) is reduced in C57BL/6J mice following 3-months of diet-induced vitamin D deficiency. These effects are not mediated by alterations in mitochondrial protein content suggesting vitamin D deficiency directly effects the mitochondrial respiratory machinery. Similar to others, we observed minimal differences in body composition following 3-months of diet-induced vitamin D deficiency in male C57BL/6J mice [19 37]. Finally our data provides evidence that vitamin D status is an important determinant of skeletal muscle mitochondrial function *in vivo* thereby supporting previous *in vitro* observations.

## Author Contributions

S.P.A and A.P conceived and designed research. S.P.A, G.F, A.M.P performed experiments and S.P.A analysed data, interpreted results and prepared figures. S.P.A, P.J.A and A.P drafted the manuscript. All authors approved the final version of the manuscript.

## References

1. Cashman KD, Dowling KG, Skrabakova Z, et al. Vitamin D deficiency in Europe: pandemic? Am J Clin Nutr 2016;103(4):1033–44 doi: 10.3945/ajcn.115.120873[published Online First: Epub Date]|.

2. Forrest KY, Stuhldreher WL. Prevalence and correlates of vitamin D deficiency in US adults. Nutr Res 2011;31(1):48–54 doi: 10.1016/j.nutres.2010.12.001[published Online First: Epub Date]|.

3. Ham AW, Lewis MD. Hypervitaminosis D Rickets: The Action of Vitamin D. British Journal of Experimental Pathology 1934;15(4):228–34

4. Bhan A, Rao AD, Rao DS. Osteomalacia as a result of vitamin D deficiency. Endocrinology and metabolism clinics of North America 2010;39(2):321-31, table of contents doi: 10.1016/j.ecl.2010.02.001[published Online First: Epub Date]|.

5. Rizzoli R, Boonen S, Brandi ML, Burlet N, Delmas P, Reginster JY. The role of calcium and vitamin D in the management of osteoporosis. Bone 2008;42(2):246–9 doi: 10.1016/j.bone.2007.10.005[published Online First: Epub Date]|.

6. Girgis CM, Clifton-Bligh RJ, Hamrick MW, Holick MF, Gunton JE. The roles of vitamin D in skeletal muscle: form, function, and metabolism. Endocrine reviews 2013;34(1):33–83 doi: 10.1210/er.2012-1012[published Online First: Epub Date]|.

7. Wicherts IS, van Schoor NM, Boeke AJ, et al. Vitamin D status predicts physical performance and its decline in older persons. J Clin Endocrinol Metab 2007;92(6):2058–65 doi: 10.1210/jc.2006-1525[published Online First: Epub Date]|.

8. Bischoff-Ferrari HA, Dietrich T, Orav EJ, et al. Higher 25-hydroxyvitamin D concentrations are associated with better lower-extremity function in both active and inactive persons aged > or =60 y. Am J Clin Nutr 2004;80(3):752–8 doi: 10.1093/ajcn/80.3.752[published Online First: Epub Date]|.

9. Houston DK, Tooze JA, Neiberg RH, et al. 25-hydroxyvitamin D status and change in physical performance and strength in older adults: the Health, Aging, and Body Composition Study. American journal of epidemiology 2012;176(11):1025–34 doi: 10.1093/aje/kws147[published Online First: Epub Date]|.

10. Beaudart C, Buckinx F, Rabenda V, et al. The effects of vitamin D on skeletal muscle strength, muscle mass, and muscle power: a systematic review and meta-analysis of randomized controlled trials. J Clin Endocrinol Metab 2014;99(11):4336–45 doi: 10.1210/jc.2014-1742[published Online First: Epub Date]|.

11. Stockton KA, Mengersen K, Paratz JD, Kandiah D, Bennell KL. Effect of vitamin D supplementation on muscle strength: a systematic review and meta-analysis. Osteoporos Int 2011;22(3):859–71 doi: 10.1007/s00198-010-1407-y[published Online First: Epub Date]|.

12. Duncan A, Talwar D, McMillan DC, Stefanowicz F, O’Reilly DS. Quantitative data on the magnitude of the systemic inflammatory response and its effect on micronutrient status based on plasma measurements. Am J Clin Nutr 2012;95(1):64–71 doi: 10.3945/ajcn.111.023812[published Online First: Epub Date]|.

13. Pleasure D, Wyszynski B, Sumner A, et al. Skeletal muscle calcium metabolism and contractile force in vitamin D-deficient chicks. The Journal of clinical investigation 1979;64(5):1157–67 doi: 10.1172/jci109569[published Online First: Epub Date]|.

14. Rodman JS, Baker T. Changes in the kinetics of muscle contraction in vitamin D-depleted rats. Kidney international 1978;13(3):189–93

15. Girgis CM, Cha KM, Houweling PJ, et al. Vitamin D Receptor Ablation and Vitamin D Deficiency Result in Reduced Grip Strength, Altered Muscle Fibers, and Increased Myostatin in Mice. Calcified tissue international 2015;97(6):602–10 doi: 10.1007/s00223-015-0054-x[published Online First: Epub Date]|.

16. Pointon JJ, Francis MJ, Smith R. Effect of vitamin D deficiency on sarcoplasmic reticulum function and troponin C concentration of rabbit skeletal muscle. Clinical science (London, England : 1979) 1979;57(3):257–63

17. Schubert L, DeLuca HF. Hypophosphatemia is responsible for skeletal muscle weakness of vitamin D deficiency. Archives of biochemistry and biophysics 2010;500(2):157–61 doi: 10.1016/j.abb.2010.05.029[published Online First: Epub Date]|.

18. McPherron AC, Lawler AM, Lee SJ. Regulation of skeletal muscle mass in mice by a new TGF-beta superfamily member. Nature 1997;387(6628):83–90 doi: 10.1038/387083a0[published Online First: Epub Date]|.

19. Seldeen KL, Pang M, Leiker MM, et al. Chronic vitamin D insufficiency impairs physical performance in C57BL/6J mice. Aging (Albany NY) 2018;10(6):1338–55 doi: 10.18632/aging.101471[published Online First: Epub Date]|.

20. Sinha A, Hollingsworth KG, Ball S, Cheetham T. Improving the vitamin D status of vitamin D deficient adults is associated with improved mitochondrial oxidative function in skeletal muscle. J Clin Endocrinol Metab 2013;98(3):E509–13 doi: 10.1210/jc.2012-3592[published Online First: Epub Date]|.

21. Bouillon R, Verstuyf A. Vitamin D, Mitochondria, and Muscle. The Journal of Clinical Endocrinology & Metabolism 2013;98(3):961–63 doi: 10.1210/jc.2013-1352[published Online First: Epub Date]|.

22. Ryan ZC, Craig TA, Folmes CD, et al. 1alpha,25-Dihydroxyvitamin D3 Regulates Mitochondrial Oxygen Consumption and Dynamics in Human Skeletal Muscle Cells. J Biol Chem 2016;291(3):1514–28 doi: 10.1074/jbc.M115.684399[published Online First: Epub Date]|.

23. Ryan ZC, Craig TA, Wang X, et al. 1alpha,25-dihydroxyvitamin D3 mitigates cancer cell mediated mitochondrial dysfunction in human skeletal muscle cells. Biochemical and biophysical research communications 2018;496(2):746–52 doi: 10.1016/j.bbrc.2018.01.092[published Online First: Epub Date]|.

24. Schnell DM, Walton RG, Vekaria HJ, et al. Vitamin D produces a perilipin 2-dependent increase in mitochondrial function in C2C12 myotubes. The Journal of nutritional biochemistry 2018;65:83–92 doi: 10.1016/j.jnutbio.2018.11.002[published Online First: Epub Date]|.

25. Romeu Montenegro K, Maron Carlessi R, Fernandes Cruzat V, Newsholme P. Effects of vitamin D on primary human skeletal muscle cell proliferation, differentiation, protein synthesis and bioenergetics. The Journal of steroid biochemistry and molecular biology 2019:105423 doi: 10.1016/j.jsbmb.2019.105423[published Online First: Epub Date]|.

26. Ashcroft SP, Bass JJ, Kazi AA, Atherton PJ, Philp A. The Vitamin D Receptor (VDR) Regulates Mitochondrial Function in C2C12 Myoblasts. American journal of physiology. Cell physiology 2020 doi: 10.1152/ajpcell.00568.2019[published Online First: Epub Date]|.

27. Health N, Council MR, Council AR. Australian Code of Practice for the Care and Use of Animals for Scientific Purposes: National Health and Medical Research Council, 2013.

28. Perry CG, Kane DA, Lin CT, et al. Inhibiting myosin-ATPase reveals a dynamic range of mitochondrial respiratory control in skeletal muscle. The Biochemical journal 2011;437(2):215–22 doi: 10.1042/bj20110366[published Online First: Epub Date]|.

29. Lanza IR, Nair KS. Mitochondrial metabolic function assessed in vivo and in vitro. Current opinion in clinical nutrition and metabolic care 2010;13(5):511–7 doi: 10.1097/MCO.0b013e32833cc93d[published Online First: Epub Date]|.

30. Picard M, Taivassalo T, Ritchie D, et al. Mitochondrial structure and function are disrupted by standard isolation methods. PloS one 2011;6(3):e18317 doi: 10.1371/journal.pone.0018317[published Online First: Epub Date]|.

31. Pesta D, Gnaiger E. High-resolution respirometry: OXPHOS protocols for human cells and permeabilized fibers from small biopsies of human muscle. Methods in molecular biology (Clifton, N.J.) 2012;810:25–58 doi: 10.1007/978-1-61779-382-0_3[published Online First: Epub Date]|.

32. Canto C, Garcia-Roves PM. High-Resolution Respirometry for Mitochondrial Characterization of Ex Vivo Mouse Tissues. Current protocols in mouse biology 2015;5(2):135–53 doi: 10.1002/9780470942390.mo140061[published Online First: Epub Date]|.

33. Miotto PM, LeBlanc PJ, Holloway GP. High-Fat Diet Causes Mitochondrial Dysfunction as a Result of Impaired ADP Sensitivity. Diabetes 2018;67(11):2199 doi: 10.2337/db18-0417[published Online First: Epub Date]|.

34. Girgis CM, Cha KM, So B, et al. Mice with myocyte deletion of vitamin D receptor have sarcopenia and impaired muscle function. Journal of cachexia, sarcopenia and muscle 2019 doi: 10.1002/jcsm.12460[published Online First: Epub Date]|.

35. Belenchia AM, Johnson SA, Kieschnick AC, Rosenfeld CS, Peterson CA. Time Course of Vitamin D Depletion and Repletion in Reproductive-age Female C57BL/6 Mice. Comparative medicine 2017;67(6):483–90

36. Visser M, Deeg DJ, Lips P. Low vitamin D and high parathyroid hormone levels as determinants of loss of muscle strength and muscle mass (sarcopenia): the Longitudinal Aging Study Amsterdam. J Clin Endocrinol Metab 2003;88(12):5766–72 doi: 10.1210/jc.2003-030604[published Online First: Epub Date]|.

37. Seldeen KL, Pang M, Rodríguez-Gonzalez M, et al. A mouse model of vitamin D insufficiency: is there a relationship between 25(OH) vitamin D levels and obesity? Nutrition & metabolism 2017;14 doi: 10.1186/s12986-017-0174-6[published Online First: Epub Date]|.

